# Identifying Essential Genes in Genome-Scale Metabolic Models of Consensus Molecular Subtypes of Colorectal Cancer

**DOI:** 10.1101/2022.10.04.510777

**Authors:** Chao-Ting Cheng, Jin-Mei Lai, Peter Mu-Hsin Chang, Yi-Ren Hong, Chi-Ying F. Huang, Feng-Sheng Wang

**Author notes:** (C.-T.C.); (F.-S.W.).

## Abstract

Identifying essential targets in genome-scale metabolic networks of cancer cells is a time-consuming process. This study proposed a fuzzy hierarchical optimization framework for identifying essential genes, metabolites and reactions. On the basis of four objectives, the framework can identify essential targets that lead to cancer cell death, and evaluate metabolic flux perturbations of normal cells due to treatment. Through fuzzy set theory, a multiobjective optimization problem was converted into a trilevel maximizing decision-making (MDM) problem. We applied nested hybrid differential evolution to solve the trilevel MDM problem to identify essential targets in the genome-scale metabolic models of five consensus molecular subtypes (CMSs) of colorectal cancers. We used various media to identify essential targets for each CMS, and discovered that most targets affected all five CMSs and that some genes belonged to a CMS-specific model. We used the experimental data for the lethality of cancer cell lines from the DepMap database to validate the identified essential genes. The results reveal that most of the identified essential genes were compatible to colorectal cancer cell lines from DepMap and that these genes could engender a high percentage of cell death when knocked out, except for EBP, LSS and SLC7A6. The identified essential genes were mostly involved in cholesterol biosynthesis, nucleotide metabolisms, and the glycerophospholipid biosynthetic pathway. The genes in the cholesterol biosynthetic pathway were also revealed to be determinable, if the medium used excluded a cholesterol uptake reaction. By contrast, the genes in the cholesterol biosynthetic pathway were non-essential, if a cholesterol uptake reaction was involved in the medium used. Furthermore, the essential gene CRLS1 was revealed as a medium-independent target for all CMSs irrespective of whether a medium involves a cholesterol uptake reaction.

**Author summary:** Essential genes are indispensable genes for cells to grow and proliferate under certain physiological condition. Identifying essential genes in genome-scale metabolic networks of cancer cells is a time-consuming process. We develop an anticancer target discovery platform for identifying essential genes that conduct cell death when the genes of cancer cells are deleted. Meanwhile, the essential genes are also inactive on their healthy cells to maintain their cell viability and smaller metabolic alterations. We use fuzzy set theory to measure metabolic deviation of the perturbation of normal cells relative to healthy and cancer templates towards predicting side effects for treatment of each identified gene. The platform can identify essential genes, metabolites and reactions for treating five consensus molecular subtypes (CMS) of colorectal cancers with using various media. We discovered that most targets affected all five CMSs and that some genes belonged to a CMS-specific model. We found that the genes in the cholesterol biosynthetic pathway are nonessential for the cells that be compensated by a cholesterol uptake reaction from a medium. Furthermore, CRLS1 was revealed as an essential gene for all CMS colorectal cancer in a medium-independent manner that is unrelated to a cholesterol uptake reaction.

## 1. Introduction

Colorectal cancer (CRC) is a worldwide health burden, it is the third- and second-ranked cancer in terms of incidence and mortality, respectively [1], indicating that the world requires improved prognoses and treatment strategies. More than 1.9 million new CRC (including anal cancer) cases and 935000 deaths were estimated to have occurred in 2020; they represent approximately one in 10 cancer cases and deaths [1]. The classification of CRC plays a pivotal role in predicting a patient’s prognosis and determining treatment strategies. The tumor, node, and metastasis (commonly abbreviated as TNM) classification system is commonly used to determine the progression of CRC; however, an in-depth characterization is necessary to improve the assessment of treatment strategies and prognoses. The consensus molecular subtype (CMS) is an RNA expression-based classification system for CRC; it was developed from 18 CRC data sets and contains 4151 CRC samples as of 2015 [2]. CRC is classified into four subtypes that each exhibit distinct molecular and biological characteristics and pathological and genetic signatures. Molecular subtype-based therapies provide a new framework for implementing preferred and precise medical treatments. Several studies have used CMS classification to predict the prognosis of a patient with CRC and determine treatment strategies for the patient [3-9].

Tissue-specific genome-scale metabolic models (GSMM) are applied to identify anticancer targets and provide detailed insights into the metabolic bases of physiological and pathological processes [10-30]. The Cancer Genome Atlas (TCGA) [31] and Human Protein Atlas (HPA) [32] databases are incorporated into human metabolic networks such as Recon X [32-37] and Human-GEM [24, 38], to reconstruct tissue-specific GSMMs for investigating metabolic behaviors. Both the TCGA and HPA databases have not used to reconstruct CMS-based tissue-specific GSMMs.

Furthermore, a literature review revealed that no study has attempted to identify targets for CRC by integrating CMS classification with genome-scale metabolic modeling. The present study used the CRC data set from TCGA to classify patient samples into five CMSs and then incorporated with Recon3D to reconstruct the CMS-specific GSMMs of CRC. We applied these CMSs to identify essential genes, metabolites, and reactions and used a fuzzy decision-making method to evaluate cancer cell mortality, healthy cell viability, and metabolic perturbation effects due to the blockage of the corresponding fluxes.

The present study proposed a fuzzy hierarchical optimization framework for identifying essential genes for treating each CMS of CRC. This framework represents an extension of the Identifying AntiCancer Targets (IACT) framework [25, 26]; it accounts for RNA-sequence expression information in inner optimization problems to yield uniform flux patterns for treated and perturbed cells. Traditional cell culture media (e.g., such as Dulbecco’s Modified Eagle Medium [DMEM] and HAM’s medium [HAM]), were designed to ensure continuous cancer cell proliferation in vitro. However, their composition do not necessarily summarize the nutritional requirements of tumor cell cultures. Studies have demonstrated that the use of specific medium components can produce cell culture results that are highly relevant to tumor metabolism [39-41]. The present study also considered various media as uptake reactions to investigate relationships of tumor cell growth with nutritional components and essential targets for CMSs of CRC.

## 2. Materials and Methods

An essential gene is an indispensable element in an organism that is essential to the survival of an organism. Because an essential gene is required for a cancer cell to grow, proliferate and survive, deleting it from a cancer cell leads to the death of this cell or causes a severe proliferation defect; by contrast, healthy cells are unlikely to be affected by its deletion. The present study developed an anticancer target discovery (ACTD) platform for identifying essential genes that can be deleted to cause the death of a cancer cell while allowing its healthy counterparts to survive with few side effects. Figure 1 illustrates the work flowchart of the ACTD platform, which mimics a wet-lab experiment that is designed for identifying essential genes.

**Figure 1.**
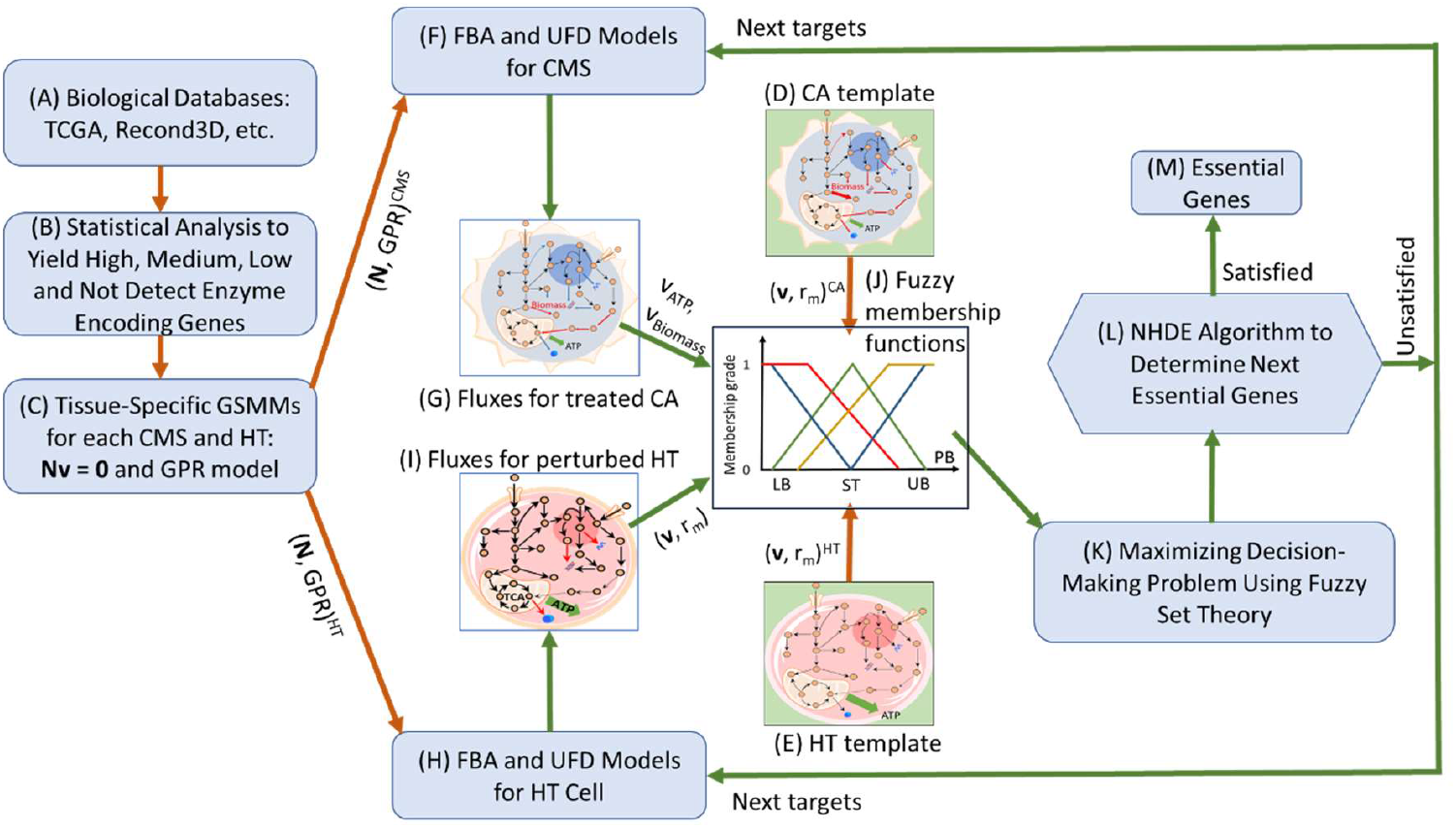
Framework for identifying essential anticancer genes. (A) Retrieval of biological data such as RNA-seq expression for cancerous cells (CA) and healthy cells (HT), and human genome-scale metabolic networks. (B) Statistical analysis of accessed RNA-seq expression data to yield high, medium, low and not detect enzyme encoding genes. (C) Reconstruction of tissue-specific genome-scale metabolic models and gene-protein-reaction models for each CMS and HT, respectively. (D) Flux distribution patterns for each CMS are obtained from clinical data (if available); otherwise a CA template can be computed through flux balance analysis (FBA) and uniform flux distribution (UFD) problems without considering dysregulated restriction. (E) Flux distribution patterns of HT cells can be obtained from clinical data (if available); otherwise, a HT template can be computed through FBA and UFD problems without considering dysregulated restriction. (F) A set of candidate genes are identified by the nest hybrid differential evolution (NHDE) algorithm and used in FBA and UFD models of CMS to compute fluxes for treatment. (G) Flux distribution and metabolite flow rates for each candidate treatment are obtained. (H) Identical genes are used for FBA and UFD models of HT cells during treatment. (I) Flux distribution and metabolite flow rates of perturbed HT cells for each candidate treatment are obtained. (J) Fuzzy membership functions for each fuzzy objective are defined to enable the evaluation of decision criterion. (K) A multiobjective objective optimization problem is converted into a maximizing decision-making problem to be solved by the NHDE algorithm. (L) Optimal essential genes are identified on the basis of decision criterion; otherwise, Steps (F) to (L) are repeated using the next set of candidate genes generated by NHDE algorithm.

### 2.1 Tissue-Specific Genome-Scale Metabolic Models

The present study combined a human metabolic network Recon3D [37] with RNA-seq expression data obtained from the TCGA database [31] to reconstruct CMS-based tissue-specific GSMMs for CRC and its healthy counterpart tissue (Figure 1A-1C). The RNA-seq expression data of 51 healthy colorectal samples (41 colon and 10 rectum samples) were downloaded from the TCGA database. A total 478 colon adenocarcinoma and 166 rectum adenocarcinoma samples were obtained as cancer samples. On the basis of CMS classification [2], the cancer samples were categorized into four subtypes that each exhibit distinct molecular and biological characteristics, namely CMS1 (microsatellite instability immune), CMS2 (canonical), CMS3 (metabolic), and CMS4 (mesenchymal). In addition, samples that could not be classified into one of the four aforementioned subtypes were categorized under CMS5 (unknown). Table 1 lists the number of accessed CRC samples that are categorized under each CMS group.

**Table 1.** Classification of colorectal cancer samples accessed from The Cancer Genome Atlas database by consensus molecular subtype (CMS). CMS classification is based on literature [2]. A sample that cannot be classified under one of the four subtypes is classified as CMS5.

|  | CMS1 | CMS2 | CMS3 | CMS4 | CMS5 |
| --- | --- | --- | --- | --- | --- |
| **No. Samples** | 83 | 226 | 72 | 151 | 112 |

RNA-seq expressions (Table 1) were used to reconstruct CMS-specific GSMMs and healthy/basal (HT) tissue. Quantile normalization was applied to normalize the raw data from healthy and CMS samples to compute the mean, confidence interval, and coefficient of dispersion for each gene. The data were then used to evaluate supportive genes and to obtain a high differential expression of enzyme-encoding genes between each CMS and its healthy cell counterpart. Recon3D comprised 2247 enzyme-encoding genes, which were classified into four levels on the basis of their participation level (i.e., high, medium, low, and not detected). Four groups of confidence reactions (i.e., high, medium, negative, and others) were obtained through gene-protein-reaction association in Recon3D. These four groups of confidence reactions were used in the CORDA algorithm [42] to reconstruct each CMS-specific GSMM and its healthy cell counterpart. Genome-scale reconstructions provide a mechanistic link between genotypes and phenotypes through stoichiometric models for metabolites and reactions. Gene-protein-reaction (GPR) associations are typically implemented as Boolean rules, which link the metabolic reactions in stoichiometric models to the gene-encoded enzymes in cells. A reaction can be catalyzed by one enzyme or isozymes. Moreover, reactions may be regulated by redundant enzymes, that is, more than two enzymes can catalyze a reactions. The redundant enzymes in a GSMM can be deleted to yield a reduced GPR association, thereby avoiding surplus computations in evolutionary optimization procedures. We developed a systems biology program that automatically builds stoichiometric and reduced GPR models in the files of the General Algebraic Modeling System (GAMS, https://www.gams.com/) for computation. The detailed procedures are discussed in another study [25].

### 2.2 Discovery of Essential Genes

The present study presents an ACTD framework to mimic a wet-lab experiment to identify essential genes (Figure 1D-1M). The framework was formulated as a fuzzy hierarchical optimization problem (Table 2).

**Table 2.** Optimization platform for anticancer target discovery design to evaluate performance of identified essential genes on basis of four fuzzy objectives.

|  |
| --- |
| **Objectives for the outer optimization problem** |
| 1. The first objective is to measure the mortality of cancer cells following gene knockout or deletion. |
| 2. The second objective is to maximize the cell viability of perturbed normal cells during gene knockout or deletion. |
| 3. The third objective is to measure the metabolic deviation of the perturbation of normal cells as a large dissimilarity relative to the cancer template. |
| 4. The fourth objective is to measure the metabolic deviation of perturbation as a close similarity relative to the normal template. |
| **subject to constraint-based models in inner optimization problems** |
| 1. FBA and UFD problems for treating cancer cells |
| 2. FBA and UFD problems for perturbation of normal cells due to treatment |

The ACTD platform was directly used to compute fluxes and metabolite flow rates for evaluating the fuzzy objectives. This method differs to that of the IACT framework, which examines logarithmic fold changes between dysregulated and template levels. An evaluation performed using the IACT framework may produce numerical inaccuracies if an evaluated value is close to zero. Furthermore, in the ACTD platform, the goal targeting cancerous (CA) cell mortality is equivalent to the minimization of both cell growth rate and adenosine triphosphate (ATP) production rate. The four goals are as follows:

The first goal is to evaluate fuzzy minimization of the cell growth rate and ATP production rate of treated CA cells as the following equations:

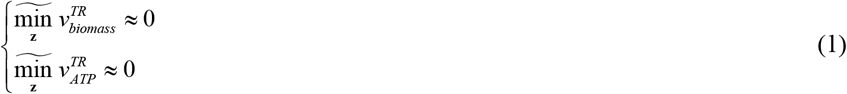

The second goal is to evaluate fuzzy minimization of the cell growth rate and fuzzy maximization of the ATP production rate of perturbed HT cells as the following equations:

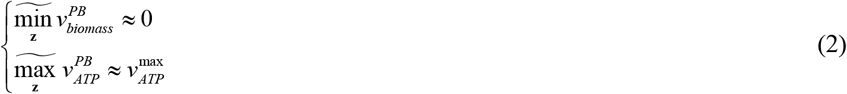

The third goal is to measure fuzzy dissimilarity of fluxes and metabolite flow rates of perturbed HT cells relative to the CA template as the following equations:

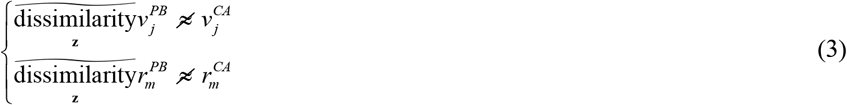

The fourth goal is to measure fuzzy similarity of fluxes and metabolite flow rates of perturbed HT cells relative to the HT template as the following equations:

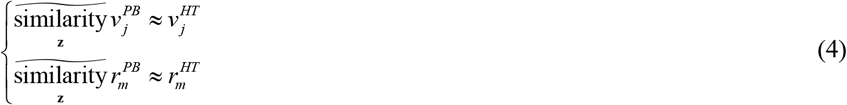

In the aforementioned equations, the decision variable **z** represents the gene encoding enzymes as determined by a nest hybrid differential evolution (NHDE) algorithm for knockout. Fuzzy minimization (i.e., 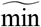) is used to evaluate the minimum cell growth rate and ATP production rate of treated CA cells. By contrast, fuzzy maximization (i.e., 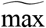) is used to meassure the maximum ATP production rate of perturbed HT cells. Fuzzy dissimilarity (i.e., 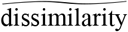) is used to evaluate the disparity of the fluxes (i.e., 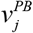) and metabolite flow rates (i.e., 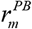) for the perturbed HT cells relative to those of the CA template. A substantial disparity means that the flux changes of the perturbed cells are considerably different from those of CA template, indicating that the perturbation of HT cells cannot lead to tumorigenesis during treatment. Fuzzy similarity (i.e., 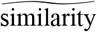) is used to evaluate the metabolic deviation between the perturbed HT cells and their template. The flow rate of the *m*^*th*^ metabolite is computed using the following equations:

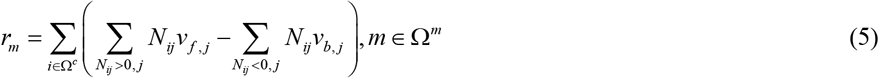

where Ω^*c*^ is the set of species located in various compartments of the cell, and *N*_*ij*_ is a stoichiometric coefficient of the *i*^*th*^ metabolite in the *j*^*th*^ reaction of a GSMM. The forward flux *v*_*f,j*_ and backward flux *v*_*b,j*_ of the *j*^*th*^ reaction are calculated by applying FBA and UFD models in the inner optimization problem as follows:

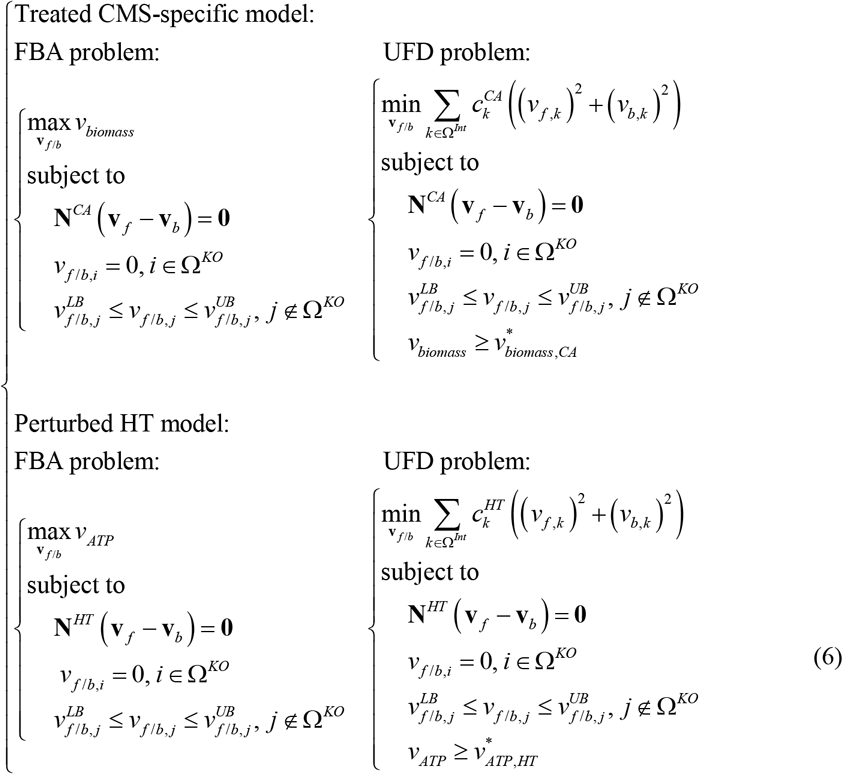

In the aforementioned equations, **N**^CA^ and **N**^HT^ are the stoichiometric matrices for each CMS-specific model and HT model, respectively, and they are reconstructed from the models presented in the previous subsection; 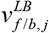 and 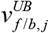 are the positive lower bound (LB) and positive upper bound (UB) of the *j*^*th*^ forward flux and *j*^*th*^ backward flux, respectively. The RNA-seq expressions for cancer and normal cells and GPR associations in Recon3D are not only used to reconstruct GSMMs but also to set the weighting factors 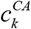 and 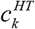 for UFD problems; the four groups of confidence reactions are assigned as follows:

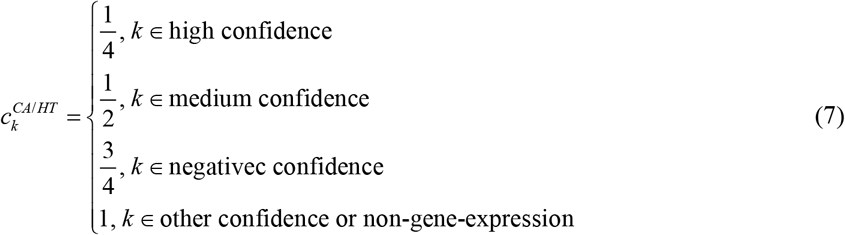

For a high confidence reaction, the smallest weighting factor is set to obtain a higher flux value from the UFD problem. Forward and backward fluxes are set to be zero if their corresponding gene encoding enzymes are knocked out. A reaction may be catalyzed by isozymes, indicating that it is still active. GPR association is used to set up the knockout reactions as follows:

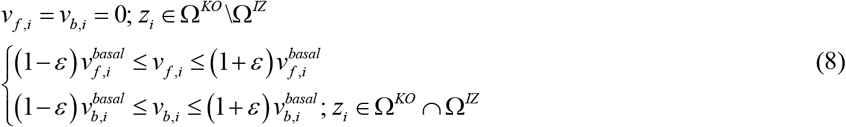

where Ω^*IZ*^ is a set of reactions regulated by isozymes that are represented in the GPR model.

### 2.3 Maximizing Decision-Making Problem

The ACTD problem expressed in Eq. (1) to (8) is a hierarchical multiobjective optimization problem (MOOP). Numerous methods can be applied for solving hierarchical MOOPs and obtain a Pareto optimal solution [25, 43, 44]. These methods are classified into two categories, namely generating methods and preference-based methods, and each method has its advantages and disadvantages [43, 44]. Generating methods apply a scalarization approach to convert a hierarchical MOOP into a single-objective optimization problem with multiple weighting factors to identify a Pareto optimal solution. By contrast, preference-based methods require a decision maker to indicate their preferences in advance before they can identify a satisfactory solution. The present study used CA and HT templates (Figure 1D and 1E) as preferences to convert an ACTD problem into a maximizing decision-making (MDM) problem through the application of fuzzy set theory (Figure 1J and 1K). One-side linear membership functions are applied to attribute fuzzy minimization (red line in Figure 1J) and fuzzy maximization (brown line) by applying the equations as follows:

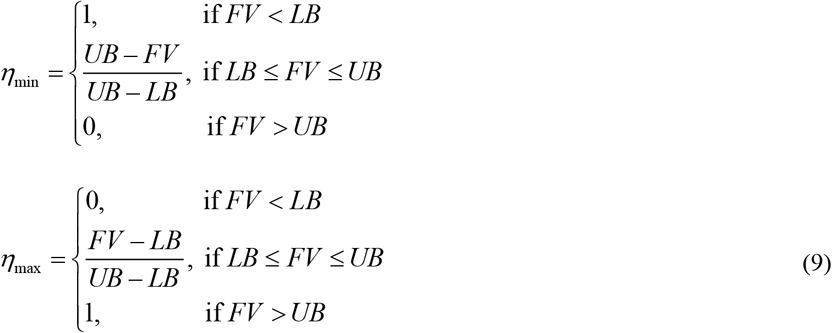

where *FV* represents the flux values computed using the treated CA or perturbed HT model. The lower bound (*LB*) and upper bound (*UB*) are obtained using the corresponding CA and HT templates, (i.e., *LB* = *ST*/4 and *UB* = 4*ST*; *ST* is the standard value for CA or HT template) used in the present study. Two-sided linear membership functions are used to attribute fuzzy dissimilarity (blue line in Figure 1J) and fuzzy similarity (green line). Fuzzy dissimilarity is a complement of fuzzy similarity; therefore, fuzzy similarity grade is derived using the equation as follows:

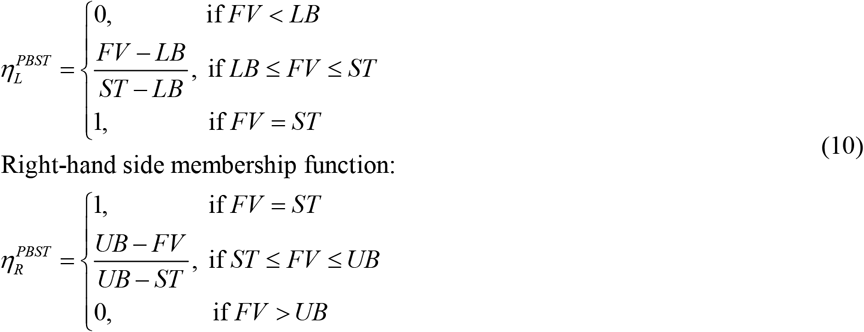

Therefore, fuzzy similarity grade is evaluated by applying the equation 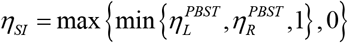, and its complement of fuzzy dissimilarity is obtained by applying the equation *ηDS* = 1 −*ηSI*.

The ACTD problem in Eq. (1) to (8) can be transformed into an MDM problem.

By applying the membership functions as follows:

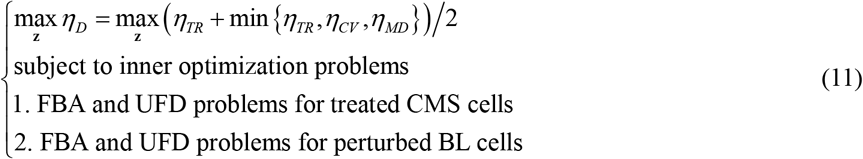

where the decision objective *η*_*D*_ is a hierarchical criterion, and the cell mortality grade *η*_*TR*_ is used to evaluate the first goal in Eq. (1) of the outer optimization problem and denoted as the first priority in the fuzzy decision-making problem. The cell viability grade *η*_*CV*_ is applied to measure the second goal in Eq. (2), and the metabolic deviation grade *η*_*MD*_ is used to measure the third and fourth goals to evaluate the mean-min level (i.e. *η*_*MD*_ = ((*η*_*DS*_ + *η*_*SI*_)/2 + min{*η*_*DS*_, *η*_*SI*_})/2). The second priority of the decision objective is used to evaluate the lowest grade in the set {*η*_*TR*_, *η*_*CV*_, *η*_*MD*_} when the cell viability or metabolic deviation grade is less than the cell mortality grade. The MDM problem in Eq. (1) is a mixed-integer optimization problem with linear and quadratic programming problems in its inner loop. It is a high-dimensional nondeterministic-polynomial-time-hard problem that cannot be solved by currently available commercial software. We employed the NHDE algorithm to solve the MDM problem. The NHDE algorithm is a parallel direct search procedure, which is an extended version based on hybrid differential evolution [45]. The detailed computational procedures are provided in the supporting information (see S1 Text). The source programs of the ACTD platform for identifying anticancer genes, metabolites and reactions of CMSs are available in http://doi.org/10.5281/zenodo.7136561.

## 3. Results

### 3.1 CMS-specific Metabolic Models

We used RNA-seq expressions for each CMS and its HT tissue to reconstruct the corresponding CMS-specific and healthy GSMMs (Figure 1A-1C). Figure 2 illustrates the numbers of species, reactions, genes, and feasible encoded enzymes for each reconstructed model. As indicated by the blue regions in Figure 2, five CMSs and HT models share numerous similarities in terms of their species, reactions, genes and enzymes. The orange regions indicate additional shared items in the five CMSs, and the grey regions illustrates the items that are specific to each CMS and HT model. Figure 3 illustrates the top 10 species and reaction classifications for each CMS and HT model. Most species in the fatty acyl groups are shared by the CMS and HT models. High percentages of organooxygen compounds, carboxylic acids, and steroids are also shared by the CMS and HT models (Figure 3A). The species that are specific to CMS2 to CMS5 comprise numerous steroid derivatives. Furthermore, extracellular transport reactions accounted for the highest percentage of reactions that occur in CMS and HT models. More than 800 fatty acid oxidation reactions are shared by CMS and HT models (Figure 3B).

**Figure 2.**
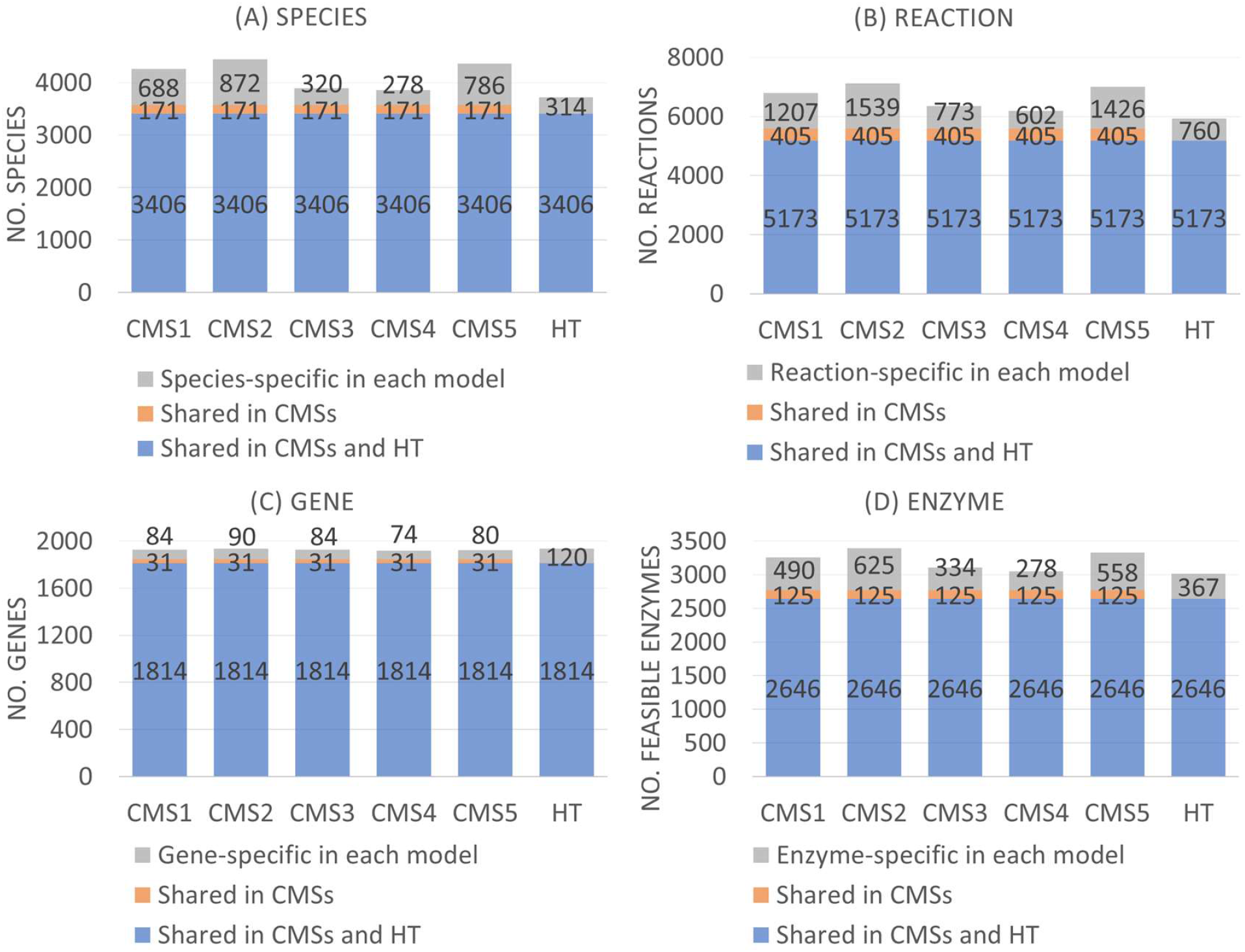
Number of species, reactions, genes, and feasible enzymes for each consensus molecular subtype (CMS) and its basal model.

**Figure 3.**
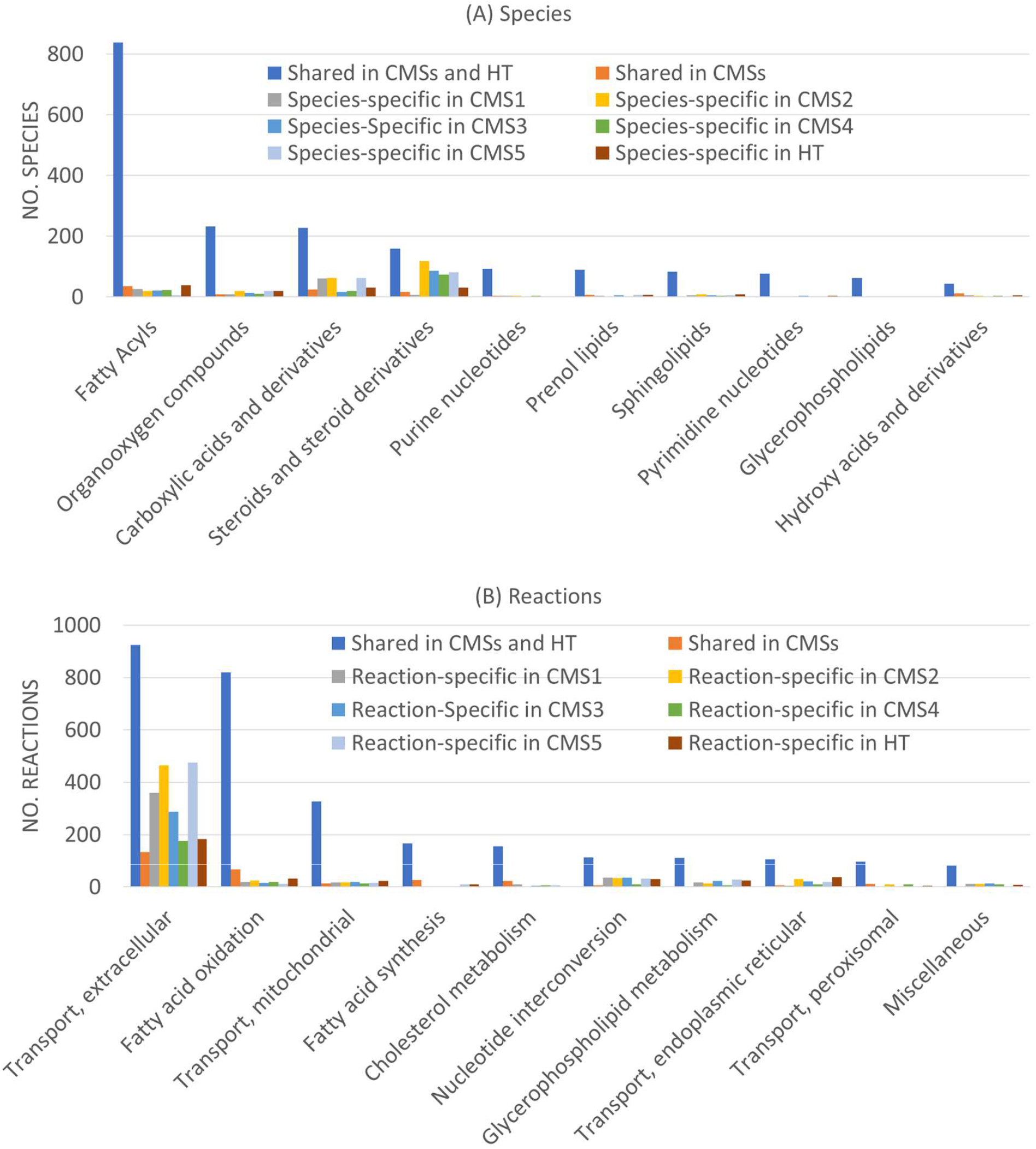
Classification of species and reactions in metabolic models of consensus molecular subtypes and their healthy cells.

### 3.2 Identifying Single Essential Genes

We first used DMEM (see S1 Table) as an uptake nutrient, and 51 uptake reactions were used to set as reversible exchangeable reactions in the computation. By contrast, secretion reactions were set as irreversible reactions. The NHDE algorithm [25-29, 43] was used to perform a series of computations to solve the MDM problem and identify a set of essential genes for each CMS (Table 3). Through the computation, we determined 26 essential genes for CMSs (Table 3), of which 20 were shared by the CMSs and six were CMS-specific. We also used a brute-force enumerate method to determine essential genes one by one to validate our computations, and the results obtained were identical to those obtained through the NHDE algorithm. Through the use of STRING (https://string-db.org/) and GeneCards https://www.genecards.org/), the protein-protein interaction (PPI) networks encoded by the 26 genes were classified into five classes (Figure 4A). The first class comprised 12 genes that are involved in cholesterol biosynthesis. The second class comprised six genes that participate in nucleotide metabolism (specifically purine and pyrimidine metabolism) and one gene in the pentose phosphate pathway. In the third class, four genes are involved in the glycerophospholipid biosynthetic pathway, whereas the others are myo-inositol and ornithine transporters.

**Table 3.** Cell mortality grades and metabolic deviation grades for essential genes of each consensus molecular subtype, as obtained using DMEM medium. N/D is obtained by dividing number of cell deaths (N) by total number of cancerous cells (D) and used in DepMap-based test. “No. Drugs” denotes number of drugs retrieved from DrugBank that modulate each gene. “SE. Score” denotes side effect score, as calculated using average adverse events.

| Gene | Cell Mortality Grade |  |  |  |  | Metabolic Deviation Grade |  |  |  |  | N/D | No. Drugs | SE. Score |
| --- | --- | --- | --- | --- | --- | --- | --- | --- | --- | --- | --- | --- | --- |
|  | CMS1 | CMS2 | CMS3 | CMS4 | CMS5 | CMS1 | CMS2 | CMS3 | CMS4 | CMS5 |  |  |  |
| DHODH | 0.660 | 0.705 | 0.670 | 0.606 | 0.691 | 0.658 | 0.682 | 0.498 | 0.404 | 0.508 | 17/54 | 26 | 0.351 |
| CAD | 0.660 | 0.705 | 0.670 | 0.606 | 0.691 | 0.654 | 0.696 | 0.508 | 0.385 | 0.500 | 27/54 | 3 | -- |
| RPIA | 0.660 | 0.705 | 0.670 | 0.606 | 0.691 | 0.638 | 0.611 | 0.549 | 0.512 | 0.516 | 6/54 | 1 | -- |
| SQLE | 0.660 | 0.705 | 0.670 | 0.606 | 0.691 | 0.575 | 0.693 | 0.548 | 0.489 | 0.488 | 2/54 | 4 | 0.415 |
| ADSS2 | 0.660 | 0.705 | 0.670 | 0.606 | 0.691 | 0.569 | 0.680 | 0.464 | 0.499 | 0.504 | 31/54 | 2 | -- |
| EBP | 0.660 | 0.705 | 0.670 | 0.606 | 0.691 | 0.568 | 0.668 | 0.450 | 0.566 | 0.504 | 0/54 | 1 | 0.412 |
| TM7SF2 | 0.660 | 0.705 | 0.670 | 0.606 | 0.691 | 0.558 | 0.702 | 0.496 | 0.496 | 0.507 | 1/54 | -- | -- |
| MVD | 0.660 | 0.705 | 0.670 | 0.606 | 0.691 | 0.557 | 0.628 | 0.509 | 0.480 | 0.495 | 49/54 | -- | -- |
| MVK | 0.660 | 0.705 | 0.670 | 0.606 | 0.691 | 0.557 | 0.628 | 0.509 | 0.482 | 0.495 | 52/54 | 1 | -- |
| PMVK | 0.660 | 0.705 | 0.670 | 0.606 | 0.691 | 0.557 | 0.681 | 0.509 | 0.480 | 0.495 | 19/54 | -- | -- |
| LSS | 0.660 | 0.705 | 0.670 | 0.606 | 0.691 | 0.556 | 0.693 | 0.547 | 0.488 | 0.489 | 0/54 | 2 | -- |
| UMPS | 0.660 | 0.705 | 0.670 | 0.606 | 0.691 | 0.554 | 0.547 | 0.546 | 0.470 | 0.532 | 29/54 | 2 | -- |
| FDFT1 | 0.660 | 0.705 | 0.670 | 0.606 | 0.691 | 0.553 | 0.617 | 0.494 | 0.496 | 0.503 | 7/54 | 1 | -- |
| HMGCR | 0.660 | 0.705 | 0.670 | 0.606 | 0.691 | 0.543 | 0.625 | 0.469 | 0.485 | 0.544 | 52/54 | 20 | 0.413 |
| CRLS1 | 0.660 | 0.705 | 0.670 | 0.606 | 0.691 | 0.539 | 0.667 | 0.469 | 0.572 | 0.494 | 35/54 | -- | -- |
| SLC7A6 | 0.657 | 0.704 | 0.668 | 0.605 | 0.690 | 0.536 | 0.598 | 0.443 | 0.491 | 0.485 | 0/54 | 1 | -- |
| PGS1 | 0.660 | 0.705 | 0.670 | 0.606 | 0.691 | 0.488 | 0.596 | 0.482 | 0.498 | 0.501 | 52/54 | -- | -- |
| SC5D | 0.660 | 0.705 | 0.670 | 0.606 | 0.691 | 0.485 | 0.696 | 0.486 | 0.503 | 0.501 | 0/54 | -- | -- |
| ADSL | 0.660 | 0.705 | 0.670 | 0.606 | 0.691 | 0.470 | 0.701 | 0.469 | 0.502 | 0.517 | 51/54 | -- | -- |
| NSDHL | 0.660 | 0.705 | 0.670 | 0.606 | 0.691 | 0.433 | 0.588 | 0.526 | 0.489 | 0.499 | 0/54 | 1 | -- |
| SLC2A13 | 0.660 | -- | -- | 0.606 | -- | 0.448 | -- | -- | 0.528 | -- | 0/54 | -- | -- |
| CYP51A1 | 0.660 | -- | -- | -- | 0.691 | 0.435 | -- | -- | -- | 0.495 | 0/54 | 5 | 0.359 |
| CTPS1 | -- | 0.705 | -- | -- | 0.691 | -- | 0.653 | -- | -- | 0.495 | 44/54 | 1 | -- |
| SLC5A3 | -- | -- | 0.670 | -- | -- | -- | -- | 0.541 | -- | -- | 2/54 | 1 | -- |
| PTPMT1 | -- | -- | -- | 0.606 | -- | -- | -- | -- | 0.502 | -- | 28/54 | -- | -- |
| PTDSS1 | -- | -- | -- | 0.606 | -- | -- | -- | -- | 0.488 | -- | 8/54 | 1 | -- |

**Figure 4.**
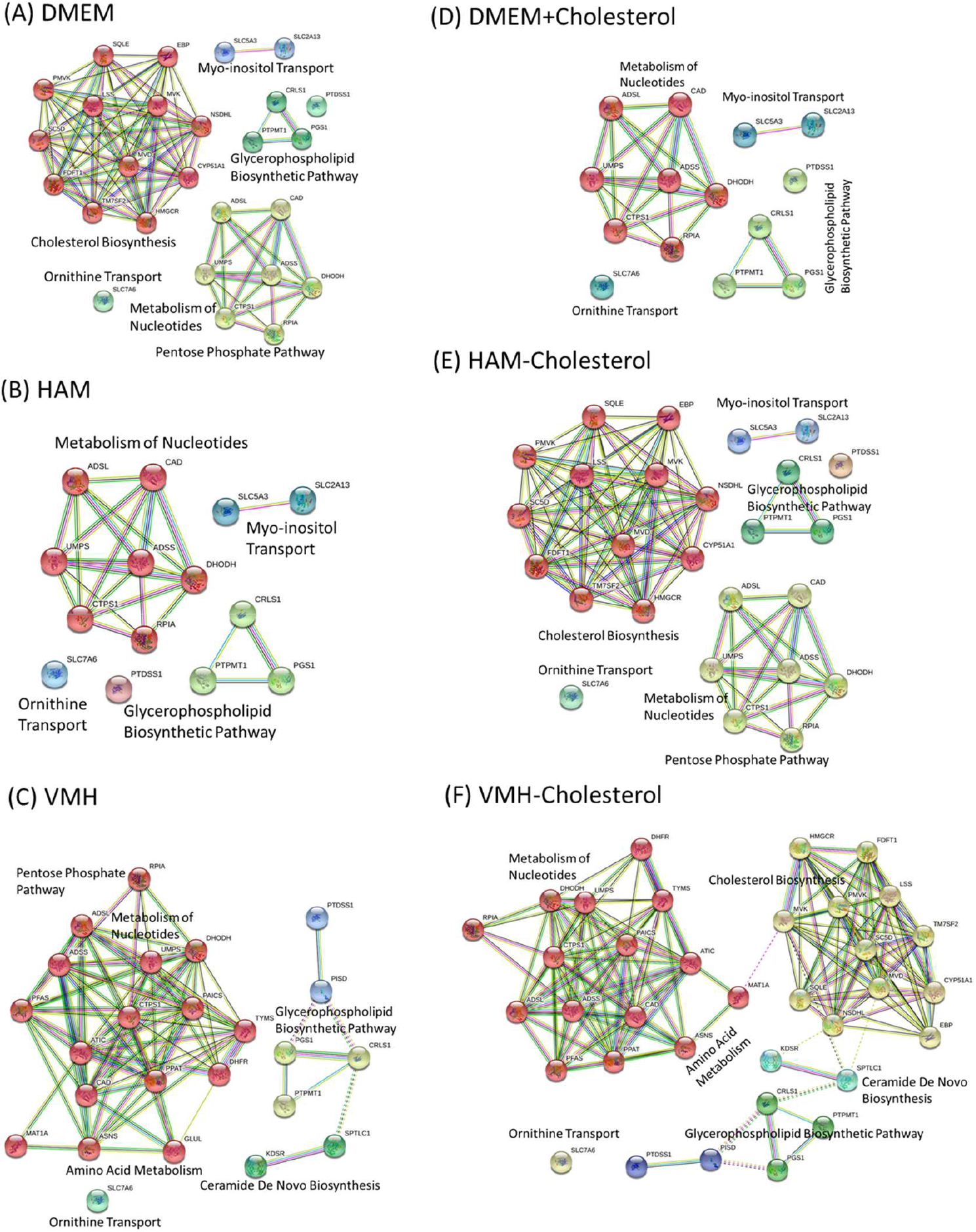
Protein–protein interactions of identified essential genes for the union set of five consensus molecular subtypes under various nutrient uptakes. (A) DMEM: Dulbecco’s Modified Eagle Medium. (B) HAM: HAM’s medium. (C) VMH: The uptake reactions are obtained from the VMH database (https://www.vmh.life/#home). (D) DMEM+Cholesterol: DMEM medium included the cholesterol uptake. (E) HAM-Cholesterol: HAM medium excluded the cholesterol uptake. (F) VMH-Cholesterol: VMH medium excluded the cholesterol uptake.

For each essential gene, the cancer cells under all the CMS classifications died (cell growth rate ≤ 10^−8^), and the ATP production rate of these cells decreased by approximately 60% relative to their maximum levels. As a result, the cell mortality grades of cancer cells was more than 0.6 (Table 2), indicating that the first fuzzy goal in Eq.(1) was achieved a 60% satisfaction level. On the basis of this computation, we assumed that the genes in HT cells were also knocked out to investigate the cell viability and metabolic deviation of perturbed cells. We discovered that perturbed HT cells maintained a cell maintenance of 10^−8^ and their maximum ATP production rate such that the second fuzzy goal was achieved with a 100% satisfaction level. The metabolic deviation grade *η*_*MD*_ was used to perform mean-min calculations to evaluate the third and fourth goals in Eq.(1), that is, for measuring the perturbation of the fluxes and metabolite flow rates of the perturbed HT cells relative to CA and HT templates, respectively. For the CMS1 model, the highest metabolic deviation grade (*η*_*MD*_) of 0.658 was achieved with the knockout of dihydroorotate dehydrogenase DHODH, and the lowest *η*_*MD*_ value of 0.433 was achieved with the knockout of sterol-4-alpha-carboxylate 3-dehydrogenase NSDHL. A higher grade indicates fewer predicted metabolic perturbation. For CMS2 to CMS5, the highest metabolic deviation grade achieved differed depending on gene that was knocked out (e.g., TM7SF2, RPIA, CRLS1 and HMGCR; Table 3).

A survey of a cancer dependency map (DepMap, https://depmap.org/portal/) revealed that most of the identified genes were compatible with the colon cancer cell lines obtained from DepMap and that the knockout of these genes (except for SLC7A6, SC5D, LSS, EBP, NSDHL and SLC2A13) resulted in a high percentage of cell death (Table 3). Some of the identified essential genes can be modulated by numerous approved drugs that are listed on DrugBank [46]. For example, 26 and 20 of the drugs surveyed from DrugBank can act on DHODH and HMGCR, respectively [46]. To investigate the grades of adverse events (AEs), the aforementioned approved drugs were used in a SIDER survey (http://sideeffects.embl.de/) involving the use of the ADDReSS (http://www.bio-add.org/ADReCS/) databases. The National Cancer Institute Common Terminology Criteria for Adverse Events (CTCAE) provide precise clinical descriptions of AE severity (from mild AE to death related to AE) grades it on a scale from 1 to 5. We used this scale to calculate the average AE (Ave.AE) and convert it into a side effect score (i.e., SE.Score = 1 – Ave.AE) that correlates with corresponding metabolic deviation grade.

### 3.3 Nutrient Uptakes

We used HAM medium as the nutrient, and 63 uptake reactions (S1 Table) were used to identify the essential genes for each CMS. The computational results (S2 Table) revealed that the cell viability and metabolic deviation grades obtained through the HAM medium were almost identical to those obtained through DMEM. We discovered that five genes participate in purine and pyrimidine metabolism, RPIA participates in the pentose phosphate pathway, PGS1 and CRLS1 participate in the glycerophospholipid biosynthetic pathway, and SLC7A6 participates in ornithine transport; these are common essential genes for all CMSs. The union set of CMSs comprised 14 identified essential genes, of which nine were shared by CMSs and five were CMS-specific. The union set was used to classify PPIs into four classes (Figure 4B), and the results revealed that the identified essential genes are not involved in the cholesterol biosynthetic pathway.

The VMH database (https://www.vmh.life/#home) published 91 uptake reactions that serve as nutrient resource for human. We used these uptake reactions (referred to as VMH medium) to investigate effects of nutrients on essential genes. Some of the uptake reactions do not exist in the reconstructed GSMMs of CMSs; thus, they were excluded in computations, and 83 uptake reactions (S1 Table) were used in computations. The NHDE algorithm revealed that 15 essential genes were shared by the CMSs and nine genes were CMS-specific (S2 Table). The union set of 24 essential genes for all CMSs was used to classify PPIs into four classes (Figure 4C), and the results revealed that the identified essential genes are not involved in the cholesterol biosynthetic pathway. The aforementioned findings indicate that these genes are nonessential on a HAM or VMH medium because the cancer cells under all CMSs will still survive if genes involved in cholesterol biosynthesis are knocked out.

Through a comparison of the uptake reactions in DMEM, the HAM medium, and the VMH mediums, the cholesterol uptake reaction was revealed to be absent from DMEM. We used three additional media to investigate the relationship of tumor cell growth with nutrient components and essential genes. DMEM was used as the first additional medium to involve the cholesterol uptake reaction (referred to as the DMEM+Cholesterol medium). The HAM and VMH media were used as the second and third media, respectively, to exclude the cholesterol uptake reaction (denoted as the HAM-Cholesterol and VMH-Cholesterol media). The ACTD platform used each medium to identify the essential genes for each CMS. Figure 4D to 4F illustrates PPI networks for the union set of the essential genes that were identified using the three additional media. The results are presented in S2 Table. When the DMEM+Cholestrol medium was used, the essential genes in the cholesterol biosynthetic pathway could not be determined (Figure 4D). However, the essential genes in the cholesterol biosynthetic pathway were identified using the HAM-Cholesterol and VMH-Cholesterol media. This finding reveals that the essential genes in the cholesterol biosynthetic pathway are determinable if the medium that is used excludes the cholesterol uptake reaction (Figure 4A, 4E and 4F). By contrast, the genes in the cholesterol biosynthetic pathway were non-essential if the cholesterol uptake reaction was involved in a medium (Figure 4B, 4C and 4D). Our simulation results are consistent with those reported by other studies [38-40], that is, nutrient components can lead to cell growth that is highly relevant to tumor metabolism. Furthermore, we also set all exchange reactions (which comprised more than 782 reactions) for each CMS as reversible uptake reactions to identify essential genes; the results indicated that only CRLS1 was determinable for all CMSs, indicating that CRLS1 was a medium-independent essential gene.

The metabolite flow distributions for CMS1 that were achieved using DMEM and the DMEM+Cholesterol medium are illustrated in Figure 5, which explains the relationship between the cholesterol uptake reaction and essential genes. Normal cells primarily metabolized glucose to pyruvate with a flux of 13.649 mmol/gDW h when DMEM was used and a flux of 13.650 mmol/gDW h when the DMEM+Cholesterol medium was used (Figure 5). Pyruvate then entered the TCA cycle to generate ATP synthesis, which is required for cell survival. The Warburg hypothesis posits that cancer cells rewire their metabolism to promote growth, survival, proliferation, and long-term maintenance. Through the CMS1 model, we discovered that CA template metabolizes large amounts of glucose to pyruvate (23.059 for DMEM and 23.046 for DMEM+Cholesterol), and which was then mostly converted to lactic acid (7.625 for both media). Thereafter, glutamine was replenished in the cycle to meet the proliferation requirement. These results are consistent with the Warburg hypothesis [47-49]. We blocked HMGCR and other genes in the cholesterol biosynthetic pathway (Figure 5) to compute specific metabolite flow rates (i.e., rates listed in the third row of data boxes in Figure 5); the results revealed that the metabolite flow rate of mev-R was 0.0001 for both media and that a series reaction occurred to produce cholesterol at level of 0.0002. Cholesterol production involves intracellular biosynthesis (discussed in an earlier subsection) and an extracellular uptake. The extracellular cholesterol uptake was zero when DMEM was used (Figure 5) because this uptake reaction was excluded such that the genes in the cholesterol biosynthetic pathway became essential. By contrast, when the DMEM+Cholesterol medium was used, the deletion of HMGCR eliminated intracellular biosynthesis; however, the cholesterol was still supplied at a level of 0.001 through the extracellular uptake reaction of the medium. Consequently, HMGCR became a nonessential gene when the DMEM+Cholesterol medium was used.

**Figure 5.**
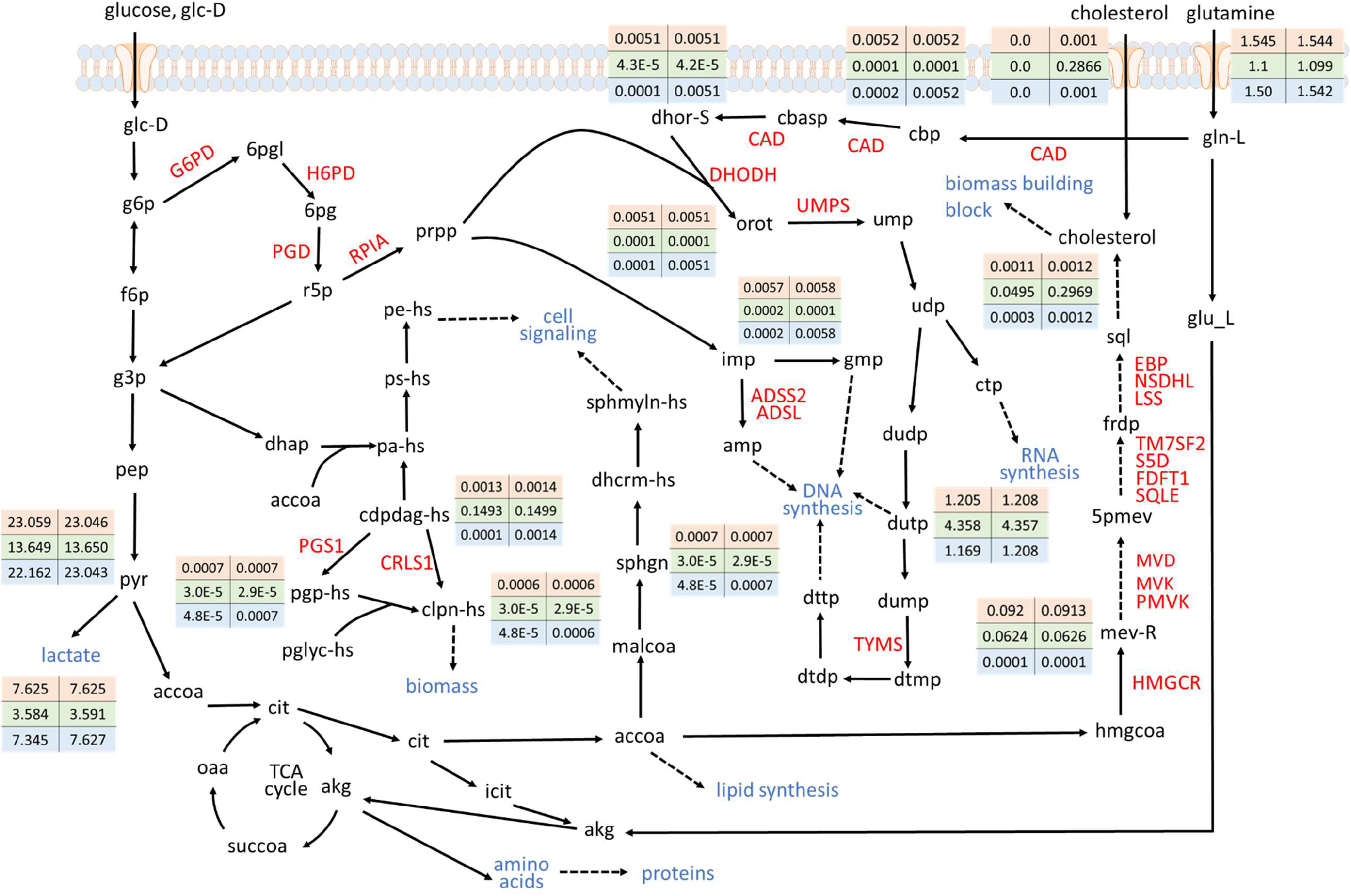
Metabolite flow rates for CMS1, as obtained using DMEM (first column in matrix of data) and DMEM+Cholesterol (2nd column) media. First column in each data box presents the computational results, as obtained using DMEM; second column presents results, as obtained using DMEM+Cholesterol medium. First and second rows of each data box present CA and HT templates, respectively; third row presents knock-out of genes in cholesterol biosynthetic pathway (e.g., HMGCR and PMVK).

### 3.4 Combination of Essential Genes

An enumerative search is a time-consuming process for identifying two target combinations of essential genes because it requires more than 4,800,000 combinations for each CMS. The ACTD platform can be used to reduce the computational burden associated with performing evolutionary procedures. Accordingly, we applied the ACTD platform by using two groups of candidates with the NHDE algorithm to identify gene combinations. The first candidate group comprised the union set of essential genes shared by CMSs (Table 1 and S2 Table), and the second group comprised the other candidate genes from feasible encoding enzymes. This strategy substantially reduced the computational time and search space to approximately 86,000 possible combinations for the two candidate groups. We performed a series of computations to obtain a set comprising two identified target combinations for various nutrient uptakes (S3 Table). On the basis of our computational results, we discovered that the metabolic deviation grades for most of the two-target combinations were superior to those for their corresponding one-target essential genes and that each combination involved at least one essential gene. Moreover, on the one-target results presented in Figure 4 indicate that a medium involving cholesterol uptake cannot be used to identify a one-target essential gene in the cholesterol biosynthetic pathway; however, a one-target essential gene can be combined with another essential gene to improve their metabolic deviation grade.

### 3.5 Essential Metabolites and Reactions

Essential metabolites and reactions denote that cancer cells that will die and the normal cells that will survive if all synthesis reactions to these metabolites are blocked. We also applied metabolite- and reaction-centric approaches on the ACTD platform to identify essential metabolites and reactions, respectively, for various media (see S4 Table and S5 Table). We compared our computational results by examining various media with or without the cholesterol uptake reaction, and we discovered that most of the essential metabolites were shared by CMSs irrespective of the presence or absence of the cholesterol uptake reaction. The computational results also revealed that the essential metabolites, farnesyl diphosphate (abbreviated as frdp in Figure 5) and formate, can be identified on a medium without the cholesterol uptake reaction, and the only exception was the CMS5 model that used VMH-Cholesterol medium. Farnesyl diphosphate is classified as a chemical compound in prenol lipids that participate in the mevalonate metabolic pathway, and it is catalyzed by FDFT1 to form squalene (sql) and to progressively synthesize cholesterol (Figure 5). Formate is an intermediate metabolite in one-carbon metabolism, and it acts as the sources and sinks in the context of mammalian cell metabolism to link various biosynthetic pathways [50]. For example, formate is released during the synthesis of cholesterol and downstream of the lanosterol step. In the present study, the deletion of formate production reduced intracellular cholesterol synthesis such that the growth of cancer cells was eliminated.

The essential reactions for all CMSs as identified using various media as listed in S5 Table. The computational results revealed that an essential reaction in the cholesterol biosynthetic pathway could not be identified irrespective of whether a medium involved or excluded the cholesterol uptake reaction. This finding differs from that obtained through a gene-centric approach (Table 3), because each essential gene that is involved in the cholesterol biosynthetic pathway through GPR association can regulate at least two reactions. We used two reaction combinations catalyzed by HMGCR to compute cell mortality and metabolic deviation grades, and the results were consistent with those obtained through a HMGCR knockout. Table 4 reveals that the reactions R_DHORD9, R_ADSL1 and R_ADSS were regulated by their corresponding genes, namely DHODH, ADSL, and ADSS1, respectively; consequently, the block of each reaction was identical to the corresponding gene knockout (Table 4). The essential gene CAD (Table 3) catalyzed three sequential reactions, namely R_CBPS, R_ASPCTr, and R_DHORTS. Therefore, the block of each reaction could eliminate the cell growth and proliferation of each CMS and achieve a satisfactory metabolic deviation grade (Table 4). The essential gene UMPS regulated two sequential reactions, namely R_OMPDC and R_ORPT, in the pyrimidine metabolic pathway such that similar results were yielded.

**Table 4.** Cell mortality grades and metabolic deviation grades for essential reactions of each consensus molecular subtype, as obtained using DMEM. Setting flux value of an essential reaction to zero leads to termination of cancer cell growth and proliferation.

| ID | Cell Mortality Grade |  |  |  |  | Metabolic Deviation Grade |  |  |  |  | Name |
| --- | --- | --- | --- | --- | --- | --- | --- | --- | --- | --- | --- |
|  | CMS1 | CMS2 | CMS3 | CMS4 | CMS5 | CMS1 | CMS2 | CMS3 | CMS4 | CMS5 |  |
| R_ASPCTr | 0.660 | 0.705 | 0.670 | 0.606 | 0.691 | 0.663 | 0.632 | 0.520 | 0.504 | 0.532 | Aspartate Carbamoyltransferase |
| R_DHORD9 | 0.660 | 0.705 | 0.670 | 0.606 | 0.691 | 0.658 | 0.682 | 0.498 | 0.404 | 0.508 | Dihydroorotic Acid Dehydrogenase |
| R_OMPDC | 0.660 | 0.705 | 0.670 | 0.606 | 0.691 | 0.655 | 0.555 | 0.530 | 0.499 | 0.526 | Orotidine-5'-Phosphate Decarboxylase |
| R_ORPT | 0.660 | 0.705 | 0.670 | 0.606 | 0.691 | 0.618 | 0.687 | 0.524 | 0.496 | 0.532 | Orotate Phosphoribosyltransferase |
| R_CBPS | 0.660 | 0.705 | 0.670 | 0.606 | 0.691 | 0.594 | 0.527 | 0.422 | 0.513 | 0.500 | Carbamoyl-Phosphate Synthase |
| R_ADSL1 | 0.660 | 0.705 | 0.670 | 0.606 | 0.691 | 0.569 | 0.680 | 0.464 | 0.499 | 0.504 | Adenylosuccinate Lyase |
| R_ADSS | 0.660 | 0.705 | 0.670 | 0.606 | 0.691 | 0.569 | 0.680 | 0.464 | 0.499 | 0.504 | Adenylosuccinate Synthase |
| R_DHORTS | 0.660 | 0.705 | 0.670 | 0.606 | 0.691 | 0.552 | 0.551 | 0.497 | 0.513 | 0.520 | Dihydroorotase |
| R_DATPtn | 0.660 | 0.705 | 0.670 | 0.606 | 0.691 | 0.504 | 0.681 | 0.498 | 0.528 | 0.547 | DATP Diffusion in Nucleus |
| R_DGTPtn | 0.660 | 0.705 | 0.670 | 0.606 | 0.691 | 0.459 | 0.702 | 0.532 | 0.574 | 0.493 | DGTP Diffusion in Nucleus |
| R_DSAT | 0.660 | 0.705 | 0.670 | 0.606 | 0.691 | 0.449 | 0.677 | 0.452 | 0.503 | 0.546 | Dihydrosphingosine N-Acyltransferase |
| R_INSTt2r | 0.660 | -- | -- | 0.606 | -- | 0.448 | -- | -- | 0.528 | -- | Transport of Inositol via Proton Symport |
| R_TRDR | 0.660 | -- | 0.670 | -- | -- | 0.510 | -- | 0.545 | -- | -- | Thioredoxin Reductase (NADPH) |
| R_INSTt4 | -- | -- | 0.670 | -- | -- | -- | -- | 0.534 | -- | -- | Transport of Inositol via Sodium Symport |
| R_PEt | -- | -- | 0.510 | -- | 0.560 | -- | -- | 0.512 | -- | 0.477 | Phosphatidylethanolamine Transport |
| R_PGPP_hs | -- | -- | -- | 0.606 | -- | -- | -- | -- | 0.502 | -- | Phosphatidylglycerol Phosphate Phosphatase |
| R_PSSA1_hs | -- | -- | -- | 0.606 | -- | -- | -- | -- | 0.488 | -- | Phosphatidylserine Synthase |

## 4. Discussion

CRC is a major disease burden, and the world requires improved prognoses and treatment strategies. Molecular subtype-based therapies provide a new potential framework for preferred and precise medical treatments. CMS classification uses RNA expressions of CRC to categorize patient samples into five subtypes. Several studies have used CMS classification to predict a patient’s prognosis and determine treatment strategies for CRC. By contrast, few studies have discussed the use of CMS classification to reconstruct CMS-specific GSMMs and to analyze the metabolic characteristics of these GSMMs. The present study used the RNA-seq expressions of CRC that were obtained from the TCGA database to reconstruct five CMS-specific GSMMs. We discovered that the five CMSs and HT models shared numerous similarities in terms of species, reactions, genes and enzymes. Most species in the fatty acyl groups and more than 800 fatty acid oxidation reactions were shared by the CMS and HT models.

The proposed fuzzy hierarchical optimization framework can identify essential genes, metabolites and reactions for treating each CMS of CRC. The optimization framework can identify essential targets that lead to cancer cell death and evaluate metabolic flux perturbations of normal cells due to treatment. Metabolic deviation grades and two-side fuzzy membership functions were used to measure the perturbations of fluxes and metabolite flow rates in the perturbed HT cells relative to HT and CA templates, respectively. A smaller metabolic deviation indicates fewer adverse effects. We used various media to identify essential targets for each CMS, and we discovered that most targets were shared by the five CMSs and that some genes belonged to a CMS-specific model. Furthermore, the essential genes in the cholesterol biosynthetic pathway were determinable if the medium used excluded the cholesterol uptake reaction. By contrast, the genes in the cholesterol biosynthetic pathway were nonessential if the medium used involved the cholesterol uptake reaction.

## Financial Disclosure

This research was supported by grants from the National Science and Technology Council, Taiwan (https://www.nstc.gov.tw/)

MOST111-2320-B-194-003 and MOST111-2221-E-194-004 to FSW

MOST111-2320-B-030-008 to JML

MOST111-2320-B-075-009 to PMHC

MOST111-2320-B-037-028;KMU-DK(A)111006 to YRH

MOST111-2320-B-A49-036 to CYFH

The funders had no role in study design, data collection and analysis, decision to publish, or preparation of the manuscript.

## Data and code availability

The source programs of anticancer target discovery platform and the tissue-specific CMS genome-scale metabolic models are coded by the General Algebraic Modeling System (GAMS, https://www.gams.com/), and are available in http://doi.org/10.5281/zenodo.7136561 Supplementary tables in Microsoft Excel format are available along with this article.

## Author contributions

Conceptualization: Jin-Mei Lai, Peter Mu-Hsin Chang, Yi-Ren Hong, Chi-Ying F. Huang, Feng-Sheng Wang

Data curation: Chao-Ting Cheng, Feng-Sheng Wang

Formal analysis: Chi-Ying F. Huang, Feng-Sheng Wang

Funding acquisition: Jin-Mei Lai, Peter Mu-Hsin Chang, Yi-Ren Hong, Chi-Ying F. Huang, Feng-Sheng Wang

Investigation: Jin-Mei Lai, Peter Mu-Hsin Chang, Yi-Ren Hong, Chi-Ying F. Huang, Feng-Sheng Wang

Methodology: Chao-Ting Cheng, Feng-Sheng Wang

Project administration: Feng-Sheng Wang.

Resources: Chao-Ting Cheng, Feng-Sheng Wang

Software: Chao-Ting Cheng, Feng-Sheng Wang

Supervision: Jin-Mei Lai, Peter Mu-Hsin Chang, Yi-Ren Hong, Chi-Ying F. Huang, Feng-Sheng Wang

Validation: Jin-Mei Lai, Peter Mu-Hsin Chang, Yi-Ren Hong, Chi-Ying F. Huang, Feng-Sheng Wang

Visualization: Chao-Ting Cheng

Writing – original draft: Jin-Mei Lai, Peter Mu-Hsin Chang, Yi-Ren Hong, Chi-Ying F. Huang, Feng-Sheng Wang

## Supporting Information

**S1 Table. Uptake reactions for DMEM, HAM medium and VMH medium**

**S2 Table. Cell mortality grades and metabolic deviation grades for essential genes of each consensus molecular subtype, as obtained using various media**.

**S3 Table. Cell mortality grades and metabolic deviation grades for combination of essential genes of each consensus molecular subtype, as obtained using various media**.

**S4 Table. Cell mortality grades and metabolic deviation grades for essential metabolites of each consensus molecular subtype, as obtained using various media**.

**S5 Table. Cell mortality grades and metabolic deviation grades for essential reactions of each consensus molecular subtype, as obtained using various media**.

**S1 Text. Description of the nested hybrid differential evolution algorithm. The source programs for identifying essential targets are coded by the GAMS software language**.

